# scART: recognizing cell clusters and constructing trajectory from single-cell epigenomic data

**DOI:** 10.1101/2023.04.08.536108

**Authors:** Jingxin Guo, Jingyu Li, Fei Huang, Jiadong Chen, Li Shen

## Abstract

The development of single-cell assay for transposase-accessible chromatin using sequencing data (scATAC-seq) has allowed the characterization of epigenetic heterogeneity at single-cell resolution. However, the sparse and noisy nature of scATAC-seq data poses unique computational challenges. To address this, we introduce scART, a novel bioinformatics tool specifically designed for scATAC-seq data analysis. scART utilizes analytical methods highly stable for processing sparse and noisy data, such as k-nearest neighbor (KNN) imputation, Term Frequency-Inverse Document Frequency (TF-IDF) weighting scheme, and the cosine similarity metric to identify underlying cellular heterogeneity in scATAC-seq data. It accurately and robustly identifies cell identities, particularly in data with low sequencing depth, and constructs the trajectory of cellular states. As a demonstration of its utility, scART successfully reconstructed the development trajectory of the embryonic mouse forebrain and uncovered the dynamics of layer-specific neurogenesis. scART is available at GitHub.

## Overview of scART

The workflow of scART is displayed in Figure 1, and the detailed strategies are shown in Figure S1. To quantify scATAC-seq data, scART first segments the genome into uniform-sized bins and uses this set of bins to count mapped sequencing reads, creating a single-cell cell-by-bin matrix (Figure 1A). Next, scART converts the raw count matrix into a binary matrix, in which 0 or 1 indicates the absence or presence of mapped reads falling within that bin (Figure 1B). To ameliorate the data scarcity problem caused by dropouts in scATAC-seq, scART adopts k-nearest neighbor (KNN) imputation to replace the potential missing values. After data imputation, biased or uninformative bins, including those from known problematic regions of the genome, bins with extremely high or low coverage among the total cells, and bins with high GC content, are filtered out to remove noises (Figure 1C and S1A). Then, scART transforms the binary matrix by Term Frequency-Inverse Document Frequency (TF-IDF) for normalization (Figure 1D). In this step, the TF process removes biases caused by uneven sequencing coverage among cells and batch effects among experiments, and the IDF process up-weights signals from cell-specific regions. To fully utilize the whole genome-wide accessibility information of each cell for dissecting cellular heterogeneity, scART directly converts the cell-by-bin matrix into a cell-to-cell similarity matrix by cosine similarity (Figure 1E). Cosine similarity measures the angular cosine value between two cell vectors according to their co-accessible region set and is more robust to noise for sparse and high-dimensional data (Andrews and Hemberg, 2018; Sohangir and Wang, 2017). From the cosine similarity matrix, scART applies truncated singular value decomposition (SVD) to generate a low-rank matrix approximation, reducing the dimensionality of the data (Figure 1E).

**Figure 1.**
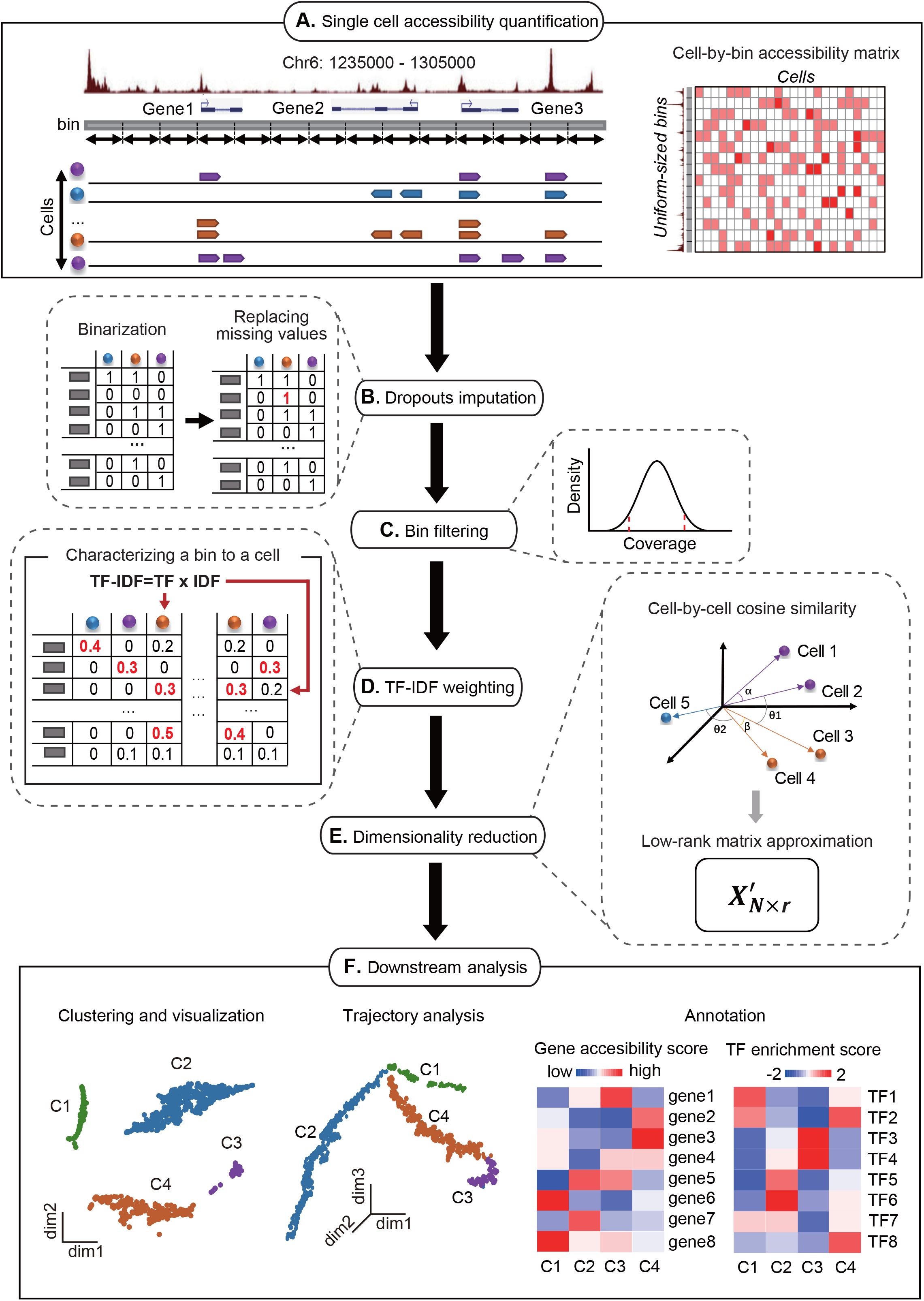
Overview of scART. (A) Single cell accessibility quantification: scART segmented the whole genome as uniformsized bins and built a chromatin accessibility matrix (Cell x Bin matrix) by counting the number of sequenced reads per cell in each of bins. (B) Dropout imputation: the raw input Cell x Bin matrix is converted to binary count matrix (with “1” indicating a specific bin is accessible in a given cell and “0” denoting inaccessible chromatin or missing data) and KNN imputation is performed to learn and fill missing values. (C) Bin filtering: undesirable bins such as low or extremely high coverage bins, bins overlapped high GC contents or black list peaks are removed. (D) TF-IDF weighting: the importance of a bin to a cell is weighted by TF-IDF weighting scheme. (E) Dimensionality reduction: converting the weighting matrix into cell-pairwise cosine similarity matrix and projecting it into lower-dimensional space by truncated SVD. (F) Downstream analysis: scART visualizes scATAC-seq data in 2-dimensional (2D) plots, builds the 3-dimensional (3D) trajectory plots and annotates clusters by cell type specific markers and motifs.

Cell clustering and trajectory analysis are performed in the lower-dimensional space (Figure 1F and S1A). For cell clustering, scART takes advantage of the density-based clustering algorithm, which depends only on the relative densities rather than a particular shape or size (Rodriguez and Laio, 2014). Trajectory analysis is performed using the discriminative dimensionality reduction tree (DDRTree) algorithm to learn the principal tree and project each cell onto its nearest location on the tree (Mao et al., 2015), and the pseudotime for each cell is assigned by the minimal spanning tree (MST) algorithm (Qi et al., 2017). Furthermore, cells within each cluster are pooled together for identification of cluster-specific accessible regions through differentially chromatin accessibility analysis (Figure 1F). Motif analysis is also carried out in each cell or group to infer cluster-specific transcription factors, which provides insights into cell type annotation (Figure 1F).

## scART identifies cell types accurately, robustly and sensitively

To benchmark scART, we compared it to six recently published algorithms for scATAC-seq analysis, including chromVAR, Cicero, LSI, cisTopic, snapATAC, and APEC. To evaluate the clustering performance of scART, we first generated a simulated scATAC-seq dataset by downsampling a previously published bulk ATAC-seq dataset of 13 human primary blood cell types that span the hematopoietic hierarchy (Corces et al., 2016), and measured each method’s performance in identifying the original cell types using the Adjusted Rand Index (ARI) (Ailon, 2008). Compared to other methods, scART most accurately clustered cells into their corresponding identities, with over 90% of cells correctly classified by scART (Figure 2A) and an ARI of 0.96 (Figure 2B). We next analyzed two real datasets generated by scATAC-seq, including a dataset of six distinct human cell lines (Buenrostro et al., 2015) and a more challenging dataset of closely related human hematopoietic cells (Buenrostro et al., 2018). For both datasets, scART resulted the highest ARI scores among the methods we tested (Figure S2A-D). While most methods were able to clearly separate distinct cell types in the dataset of six cell lines (Figure S2A and S2C), only scART achieved an ARI score above 0.5 when analyzing the dataset of human hematopoietic cells (Figure S2B and S2D). Although APEC is second only to scART in the analysis of human-hematopoietic-cell dataset (Figure S2B), it showed an unexpected low ARI in the analysis of six-cell-line dataset (Figure S2A), likely due to its susceptibility to batch effects in scATAC-seq experiments (Figure S2E). In contrast, scART exhibited its stability against batch effects (Figure S2E).

**Figure 2.**
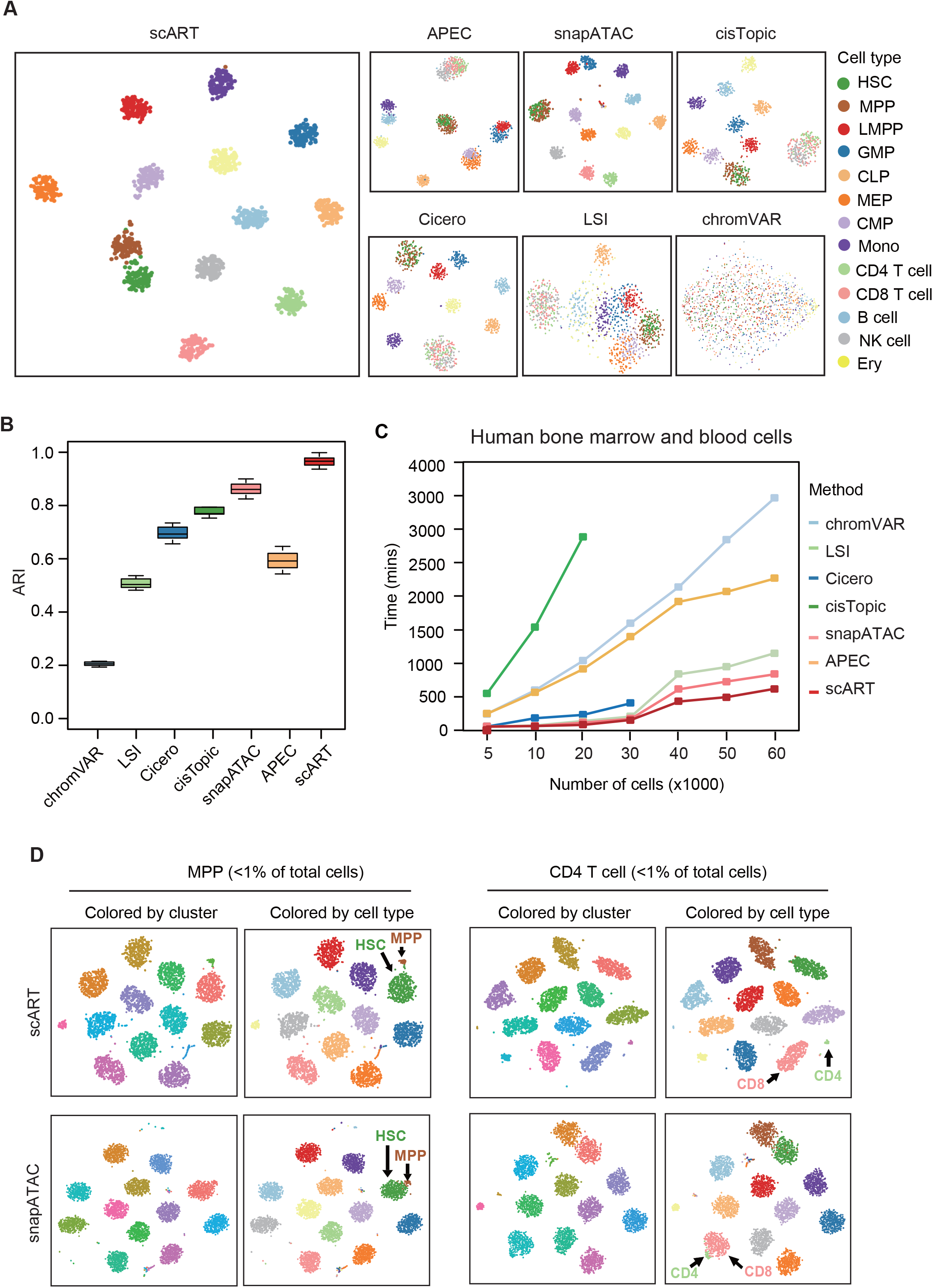
scART identifies cell types accurately, robustly and sensitively. (A) Accuracy. 2D t-Distributed Stochastic Neighbor Embedding (tSNE) diagrams of the simulated HSC scATAC-seq datasets showed the clustering results of different methods, cells are colored by cell type. The human hematopoietic cell lines contain 13 cell types including HSCs (hematopoietic stem cell), MPP (multipotent progenitor cells), CMP (common myeloid progenitor), GMP (granulocye-macrophage progenitor), LMPP (lymphoid-primed multipotential progenitors), CLP (common lymphoid progenitor), MEP (megakaryocytic-erythroid progenitor), Mono (monocyte), Ery (erythroid), B (B cells), CD4 (CD4+ T-cells), CD8 (CD8+ T cells), and NK (natural killer cells). (B) Robustness. Adjusted Rand index (ARI) showed the clustering accuracy of each method on the simulated HSC scATAC-seq datasets. Center line, median; box limits, upper and lower quartiles; whisker, 1.5 X interquartile range; outliers were removed. (C) Scalability. Visualization of scART and snapATAC on identifying rare population cells that account for less than 0.1% of the total population by simulated scATAC-seq datasets from bulk ATAC-seq data of 13 human hematopoietic cell lines. (D) Sensitivity. The computing time required for different algorithms to cluster simulated scATAC-seq with cell number from 5,000 to 100,000, sampled from the 13 previously published bulk ATAC-seq datasets.

In addition to the accuracy of cell clustering, high sensitivity to rare cell populations is also crucial in scATAC-seq data analysis. Among the published methods, only snapATAC has been demonstrated to have the ability to effectively detect rare cell populations. Therefore, we compared scART with snapATAC in identifying rare populations by using two simulated datasets that we generated from bulk ATAC-seq datasets. In the simulated datasets, MPP cells or CD4 T cells only made up less than 1% of the total cells. As shown in Figures 2C, scART successfully dissected the rare population of MPP cells from HSC cells, as well as the rare population of CD4 T cells from CD8 T cells. Taken together, these results demonstrate that scART is both accurate and sensitive in identifying cellular heterogeneity.

## scART exhibits superior performance with low sequencing depth

Although high sequencing depth is usually required to generate high complexity, unbiased, and interpretable datasets, many single-cell datasets are generated with insufficient sequencing depth (Rizzetto et al., 2017; Sims et al., 2014), which can further exacerbate the intrinsic sparsity problem of scATAC-seq data. To evaluate the performance of scART in analyzing datasets with low sequencing depth, we generated a series of simulated scATAC-seq datasets from bulk ATAC-seq datasets of human hematopoietic cell lines, ranging from 10,000 reads per cell (high coverage) to 2,500 reads per cell (low coverage). As the sequencing depth was reduced to lower than 5,000 reads per cell, all of the peak-dependent methods (i.e., chromVAR, Cicero, LSI, cisTopic, and APEC) lost the ability to identify cell types, with ARI values lower than 0.5 (Figure 3A and S3). In contrast, scART and snapATAC consistently identified cell types even in the datasets with 2,500 reads per cell (Figure S3). At low sequencing depth, scART was more sensitive in uncovering differences among closely related cell types, such as HSC and MPP (Figure 3B). We also benchmarked the sensitivity of scART, snapATAC, and cisTopic on a low-coverage scATAC-seq dataset of adult mouse brain with an average sequencing depth of 3,000 reads per cell, and found that scART outperformed the other methods in identifying the complex cellular heterogeneity in the brain (Figure 3C and S4). Collectively, these results demonstrate that scART outperforms other methods in the analysis of scATAC-seq datasets of different sequencing depth.

**Figure 3.**
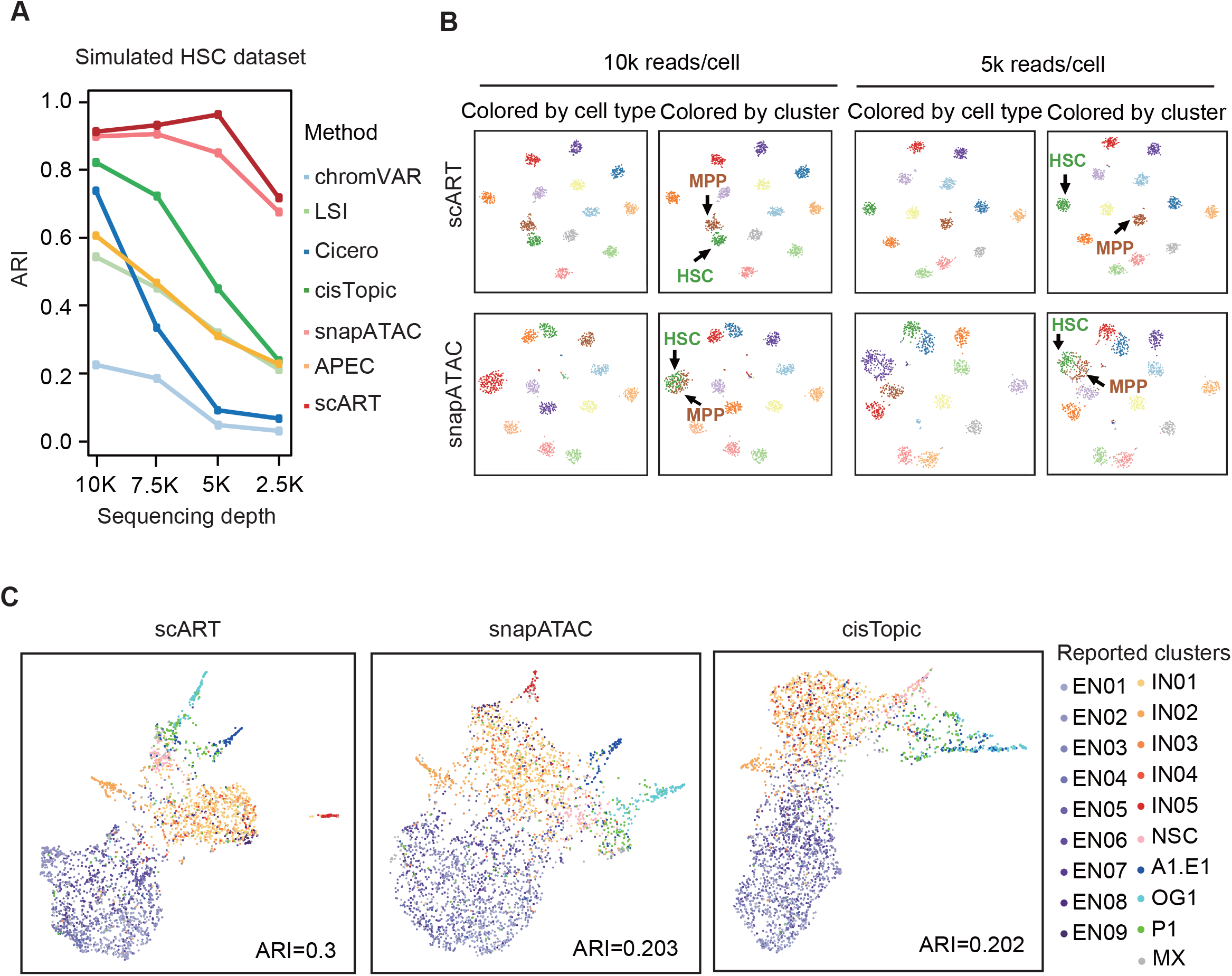
scART exhibits superior performance with low sequencing depth. (A) Visualization of scART and snapATAC on distinguishing minor chromatin accessibility variation among cells by simulated scATAC-seq dataset from bulk ATAC-seq data of 13 human hematopoietic cell lines with varying coverages. (B) 2D uniform manifold approximation and projection (UMAP) plots of adult mouse brain dataset from different methods. Cells were colored based on the reported clusters. As reported, the clusters were assigned based on both marker genes and scATAC-seq signals.

## scART reconstructs the developmental trajectory of embryonic mouse forebrain

Chromatin accessibility has been shown to predict the future expression of lineage-determining genes and thus the lineage choice of a differentiating cell (Clyde, 2021; Llorens-Bobadilla et al., 2020). Therefore, it is possible to infer cellular transitions during development based on single-cell chromatin accessibility profiles. In scART, we integrated the MST and DDRTree algorithms used in reserved graph embedding (RGE), a population graph-based pseudotime analysis algorithm in scRNA-seq analysis, to predict the developmental trajectory based on the lower dimensional space that the cells lie upon and use a cell-cell graph to describe the structure among cells. By default, scART identifies branch points that describe significant divergences in cellular states automatically.

As a demonstration, we applied scART to reconstruct the differentiation trajectory of human hematopoietic cells (Figure S5A and S5B). The trajectory successfully reflected the three main biological programs during HSC differentiation, including lymphoid, erythroid, and myeloid transcriptional programs (Figure S5B). Indeed, by evaluating the deviation of main TFs along the trajectory, we found that the accessible regions in HSC and MPP cells were enriched with HOX motif, and those in the myeloid, erythroid, and lymphoid lineages enriched with GATA1, CEBPB, and EBF1 motifs, respectively (Figure S5C and S5D) (Buenrostro et al., 2018; Schep et al., 2017; Vilagos et al., 2012).

We further demonstrated the utility of scART by applying it to infer the development trajectory of neurogenesis in embryonic forebrain development, which is more challenging as the cell dynamics can be very complex. We utilized a dataset containing 12,733 high-quality scATAC-seq profiles derived from fetal mouse forebrains at seven developmental stages (Preissl et al., 2018). The trajectory inferred by scART predicted three developmental branches: the inhibitory neuron development branch (IN branch), the excitatory neuron development branch (EX branch), and the glia cell development branch (Glia branch) (Figure 4A, 4B, and 4C). Twelve distinct cell populations were identified, including three radial glia-like cell groups (RG1-3), three excitatory neurons (EX1-3), four inhibitory neurons (IN1-4), astrocytes (AC), and erythro-myeloid progenitors (EMP). Note that the three distinct excited neurons that scART identified were mixed-clustered in the original study, and the distinct populations exhibited changes in abundance through development (Figure 4D and 4E). EX1 and EX2 were found at early developmental stages and exhibited characteristics of deeper cortical layer neurons with high gene accessible scores for deeper layer markers such as Foxp2 and Bcl11b (Figure 4B) (Chen et al., 2008; Chiu et al., 2014). In contrast, EX3 represented a group of upper cortical layer neurons with high accessible scores of upper layer markers such as Calb1 and Cux1 (Cubelos et al., 2010; Gray et al., 2017) (Figure 4B). Additionally, the EX3 cluster cells expanded at E14.5-E15.5, consistent with the timing of upper layer formation (Jabaudon, 2017; Molyneaux et al., 2007) (Figure 4D and 4E). Overall, scART accurately clustered embryonic forebrain cells and constructed the trajectory of neurogenesis and gliogenesis programs with sensitivity and precision.

**Figure 4.**
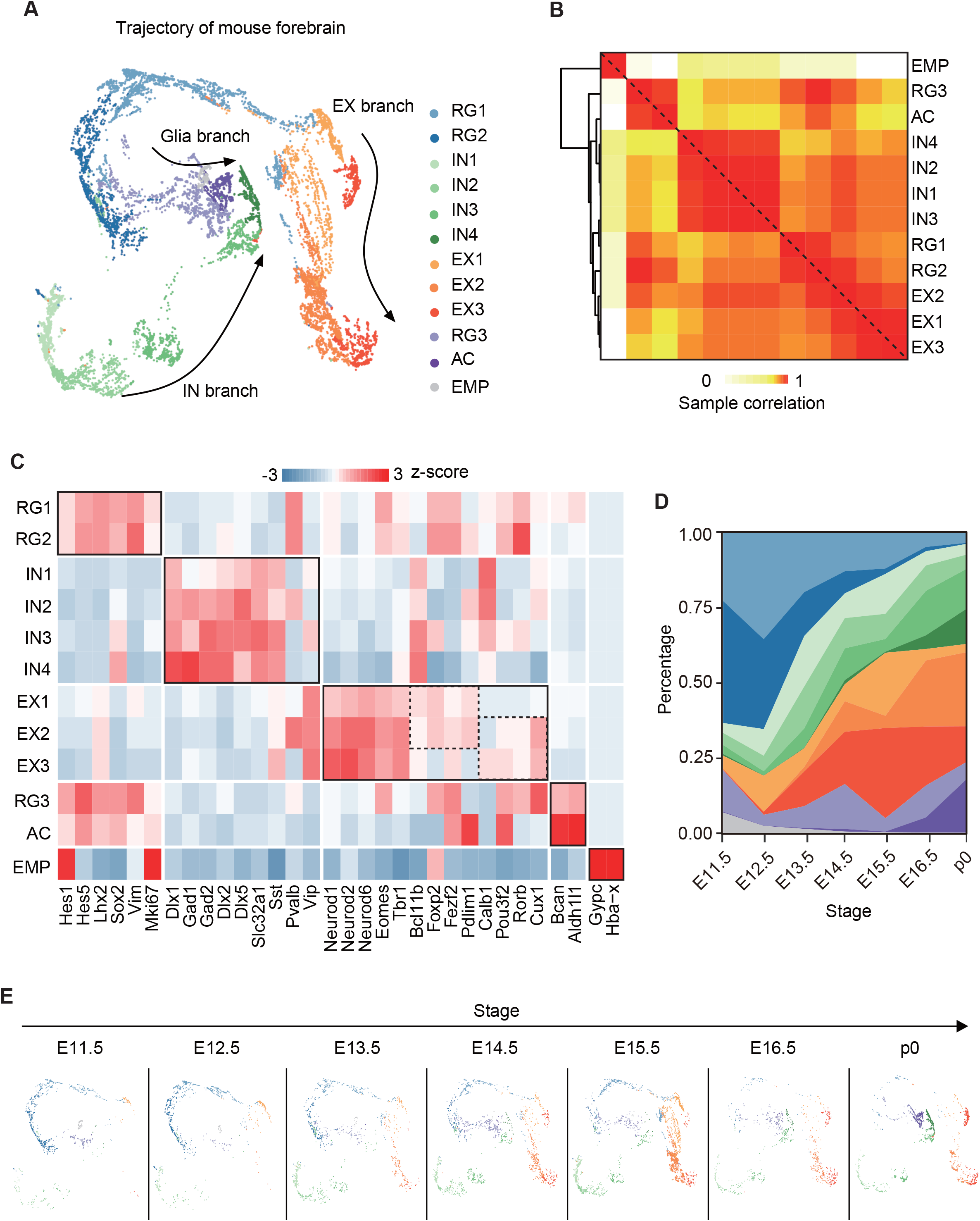
scART reconstructs the developmental trajectory of embryonic mouse forebrain. (A) 3D visualization showed the pseudotime trajectory of embryonic mouse forebrain development. scART clustered 12377 cells from 7 different developmental stage into 12 cell types including 3 group of radial glia cells (RG1, RG2 and RG3), 4 group of inhibitory neuron cells (IN1-4), 3 group of excitatory neuron cells (EX1-2), astrocyte cell (AC) and erythromyeloid progenitors cell (EMP). (B) Average scores of the marker genes for each cell cluster identified by scART. The score values were normalized by the standard score (z-score). The bottom row showed the cell types annotated according to the gene score of cell type specific markers. (C) Hierarchical clustering of the cluster-cluster correlation matrix. (D) Quantification of cellular composition was calculated at each development stage. (E) The 3D trajectory diagrams at different stage visualized the dynamic changes of annotated cell types.

## Notes

### Competing Interest Statement

The authors have declared no competing interest.

